# Development of an ELISA assay for EV quantification to assist the large-scale EV manufacture

**DOI:** 10.1101/2024.10.31.621446

**Authors:** Xin Zhou, Wenli Wang, Lei Zhang, Shiyi Yang, Xincheng Peng, Xin Zhang, Jinxiu Zhao, Xinjun He, Ke Xu

## Abstract

Accurate EV quantitation is essential for ensuring quality, consistency, and safety during large-scale extracellular vesicle (EV) manufacturing. In the upstream phase, EV quantitation allows for the monitoring of cell culture conditions that impact EV yield and quality, while in the downstream phase, it helps to control the efficiency and purity of EV purification. Nanoparticle Tracking Analysis is the most commonly used method for EV quantification, while it faces significant limitations due to interference from nanosize contaminants like protein aggregates, especially in crude samples (e.g. cell culture media). To address this issue, we developed a highly specific and accurate ELISA assay that quantifies EVs even in crude samples. With ultra-pure EV standard samples, this assay showed reliable quantitative result of EV detection to support method development as well as in-process control of large-scale EV manufacture. The detection range of this assay is from 4.1E7 to 3E10 EVs/mL, with an LOD of 1.04E7 EVs/mL and an LOQ of 3.21E7 EVs/mL. We therefore developed this assay into a testing kit and demonstrated that this EV quantification ELISA kit is capable of ensuring minimal interference from impurities and supporting the process development and in-process control in EV production.

## Background

Extracellular vesicles (EVs) are small, membranous particles released by cells into the extracellular space. EVs carry a variety of biomolecules, such as proteins, lipids, RNA species, reflecting the cell of origin.^1^ They play key roles in intercellular communication by transferring their cargo between cells, influencing physiological processes like immune responses,^2^ tissue repair,^3^ and inflammation.^4, 5^ EVs transfer functional molecules between cells, modulating processes like immune regulation and tissue regeneration.^6^ Also, EVs contribute to signaling pathways that regulate cellular growth, differentiation, and response to environmental changes.^7-10^ Therefore, EVs can help to understand disease mechanisms and the complex networks of cellular communication, serving as biomarkers for diseases such as cancer,^11-17^ cardiovascular diseases,^18-20^ and neurodegenerative diseases,^21-23^ as their content often reflects pathological changes in the body. Also, EVs are capable of working as nano-carriers for drug delivery due to their ability to naturally transfer molecules between cells.^24-29^ Hence, EVs hold great potential in broad applications in the field of clinical diagnostics and therapeutics.

Although EV exploration is rapidly expanding in academic studies, increasing production to meet industrial demands remains challenging in the development of scalable and reliable process.^30-34^ Among all the possible tasks, precise quantification of EVs is essential throughout both upstream and downstream process development. In the upstream phase, EV quantification is crucial for tracking cell culture conditions that affect EV yield and quality. During the downstream phase, it plays a key role in managing purification steps, ensuring effective EV recovery and purity. However, Nanoparticle tracking analysis (NTA), the most widely used EV quantification method,^35-38^ faces limitations in quantifying EVs in crude samples, such as cell culture media, due to the presence of various non-EV particles and contaminants.^39^ NTA works by detecting and tracking individual particles based on their Brownian motion to measure size and concentration. However, crude samples like cell culture media contain a mixture of particles, including proteins, cell debris, and other extracellular components, which can interfere with the analysis. Many other EV quantification methods available on the market also have inherent limitations. For example, while nano flow cytometry (nanoFCM) can effectively differentiate between EVs and particle contaminants, its high instrument cost and low throughput limit its application.^40-42^ Sandwich ELISA (Enzyme-Linked Immunosorbent Assay), which uses antibodies to recognize antigens on the surface of EVs, can specifically identify and quantify EVs. There are many commercial ELISA kits that were reported to be utilized in a numbers of academic research works for EV quantification, such as the ExoTEST from Hansabiomed,^43-45^ the CD9-Capture Human Exosome ELISA Kit from Wako,^46, 47^ the ExoELISA kit from SBI,^48, 49^ the Exo-Quant from BioVision,^50, 51^ etc. However, few commercial ELISA kit provide the information of the EV parent cell line as well as the purity of the EV standard sample within the kits. In addition, most of the commercial ELISA kit only provide the mass concentration of their EV standard without particle number concentration. These features make them more appropriate for academic research application rather than industry large-scale EV manufacture.

To address this issue, herein we developed an assay named Vesicure human EV (CLIA) ELISA detection kit. The kit specifically captures EVs through anti-CD81 coated on the plate, and detects them using anti-CD9 and anti-CD63, ensuring the specificity of EV detection in crude sample. In addition, it achieves sensitive EV quantification based on the principle of enzyme-catalyzed chemiluminescence. More importantly, highly purified EVs from HEK293 or MSC cells are used as the standard sample, ensuring the accuracy of the measurement.

## Methods and Material

### Vesicure total human EV quantification ELISA

The ELISA assay was performed by following the manual of the product. Briefly, all reagents and samples were equilibrated to room temperature prior to the assay. The coated plate was washed once with 300 μL/well of washing buffer and tapped dry. 100 μL/well of standard and test samples was added to each well, respectively, and incubated for 2 hours at 37°C. Wash the wells three times, then add 100 μL of the detection antibody and incubate for 1 hour at 37°C. Wash again, add 100 μL of Streptavidin-HRP, and incubate for 30 minutes at 37°C. After washing five times, 100 μL of the chemiluminescent substrate was added and incubate for 15 minutes at room temperature, protected from light. Finally, chemiluminescent signal was measured in a plate-reader.

### Density gradient ultracentrifugation (DGUC)

To purify extracellular vesicles (EVs) using density gradient ultracentrifugation, first, concentrate 2000 mL of cell supernatant by about ten-fold into 200 mL. Centrifuge at 6000 g for 20 minutes at 4°C and collect the supernatant. Treat the supernatant with 1 mM MgCl_2_ and 20 U/mL of Benzonase for 16 hours at 25°C (or 3 hours at 37°C) to digest nucleic acids. Next, centrifuge at 16,000 g for 30 minutes at 4°C, collecting the supernatant. Divide this supernatant into six tubes, fill with PBS, and balance. Centrifuge at 133,900 g for 60 minutes at 4°C and discard the supernatant. Resuspend the pellet in PBS, transferring equal volumes to new tubes. Prepare 17.5% and 45% iodixanol solutions, layering them in Quick-Seal Ultra-Clear tubes with PBS at the top. Centrifuge at 150,000 g for 16 hours at 4°C. Carefully collect the white interface layer and transfer it to new tubes, followed by centrifugation at 20,000 g for 30 minutes. Finally, resuspend the EV pellet in 100 μL of PBS and quantify using ZetaView, storing the aliquots at 2-8°C at a concentration of 1E13 particles/mL.

### TEM imaging

EVs were gently placed onto a 200-mesh copper grid coated with a carbon film, which had been glow-discharged to improve sample attachment and reduce surface hydrophobicity. For negative staining, the EVs were treated with a 2% uranyl acetate solution at room temperature for 1 minute, enhancing contrast by surrounding the vesicles with an electron-dense material. After staining, excess uranyl acetate was promptly rinsed off with distilled water to avoid excessive staining. The grids were then allowed to air-dry for stability during examination. EVs were subsequently imaged using a Tecnai G2 transmission electron microscope (Thermo FEI) at 120 kV, enabling high-resolution visualization of EV structure for detailed morphological analysis.

### Nano-Tracking Analysis (NTA)

Extracellular vesicles (EVs) suspended in PBS were diluted with ultra-pure water to a concentration around 1E8 particles/mL. The diluted samples were then directly introduced into the ZetaView Nano Particle Tracking Analyzer (ParticleMetrix, PMX120-Z) for measurement.

### Size Exclusion High-Performance Liquid Chromatography (SE-HPLC)

The SE-HPLC analysis was performed with Sepax SRT SEC-300 (5μm, 7.8×300 mm) on an Agilent 1260 Infinity II HPLC system, with a mobile phase of 100mM phosphate buffer containing 200mM NaCl, pH 7.2. Isocratic elution was employed with a flow rate setting at 1 mL/min.

## Result and Discussion

Accurate EV quantitation is critical in the process development and in-process control (IPC) of large-scale EV manufacturing to ensure product quality, consistency, and safety throughout both upstream and downstream process development. For upstream, EV quantitation is vital for monitoring the EV yield in different culture conditions. For downstream, EV quantitation helps control purification steps like chromatography, ultrafiltration, and size exclusion to ensure efficient EV recovery and purity. However, the most commonly used EV quantification method, NTA, struggles to quantify EVs in crude samples like cell culture media due to the presence of similarly sized particles such as protein aggregates and debris in crude, thus making it inappropriate for EV measurement to support upstream and downstream process development and in-process control.

To address this issue, we developed an ELISA assay and kit for the accurate quantification of human EVs, as shown in Figure 1A. The working principle of the EV quantitation ELISA kit involves capturing and detecting EVs using specific antibodies, as illustrated in Figure 1B. Anti-CD81 is coated on a 96-well plate to serve as the capture antibody, binding EVs present in the sample. Biotinylated anti-CD63 and anti-CD9 are used as detection antibodies, targeting other EV surface markers to confirm the presence of EVs. HRP-Streptavidin is then added, binding to the biotin on the detection antibodies. This enzyme catalyzes the substrate, producing a luminescent signal, which is measured to quantify the amount of EVs in the sample. In this assay, we take advantage of the antibody-antigen interaction to target three different tetraspanin EV surface proteins in order to ensure the specificity for EV measurement, making it ideal for EV detection in crude samples throughout upstream to downstream with the presence of non-EV nanoparticles (e.g. cell culture media as shown in Figure 1C).

**Figure 1.**
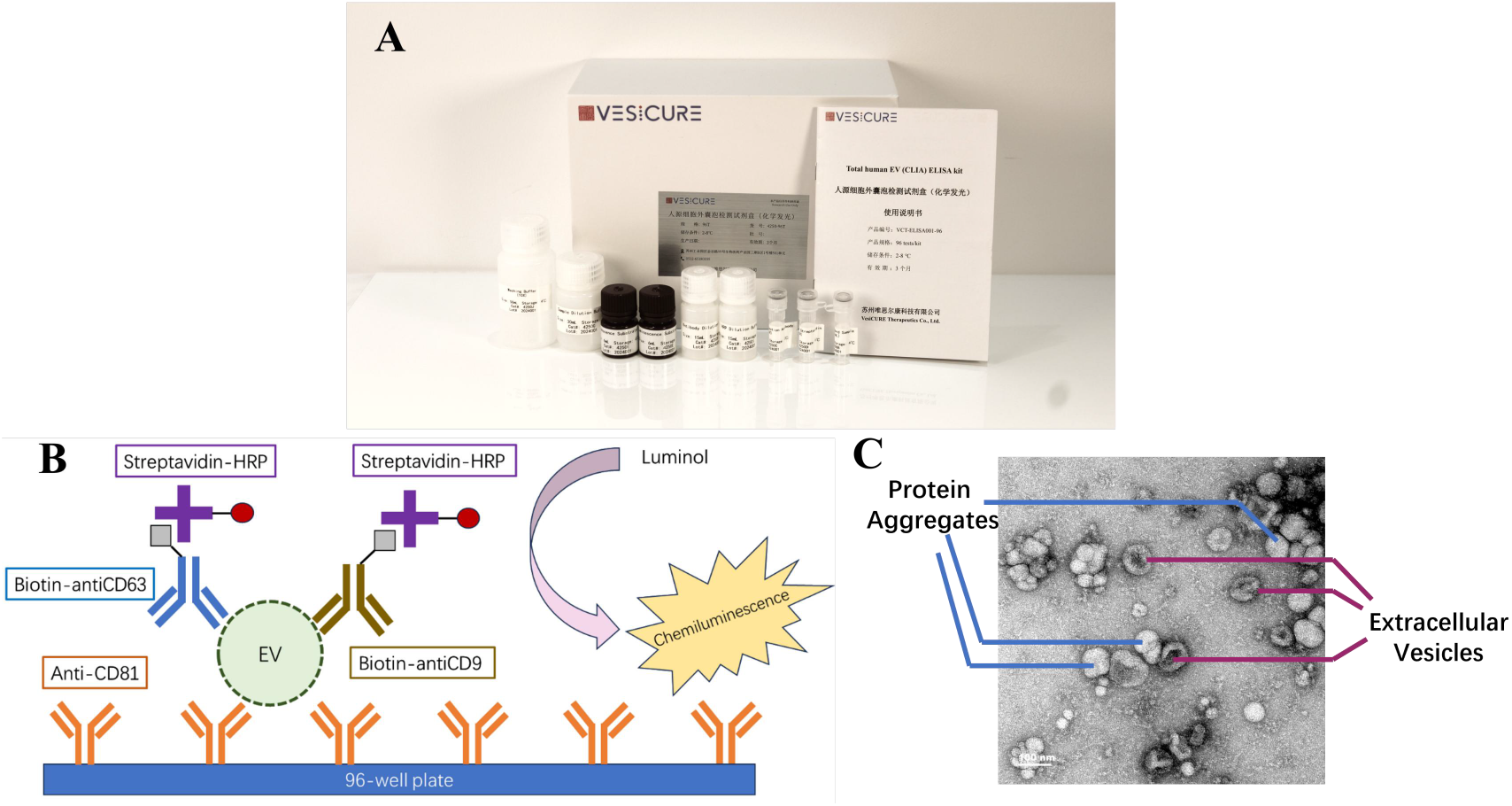
Introduction of the Vesicure total human EVs quantitation ELISA (CLIA) kit (A) Photo of the ELISA kit product (B) Schematic principle of the Vesicure total human EVs quantitation ELISA (CLIA) Assay (C) TEM image of UC-Purified EV sample from cell culture media

The purity of the EV standard sample is crucial in the EV quantification ELISA assay because it directly impacts the accuracy and reliability of the results. A pure EV standard provides an accurate NTA readout of the EV sample without the presence of contaminates, ensuring the correct EV particle concentration to calibrate the ELISA assay. To obtain the highly pure EV sample for our ELISA assay, density gradient ultracentrifugation (DGUC) was utilized on harvested cell culture fluid. DGUC is one of the most widely used method to get ultra-pure EVs based on EVs physical properties (i.e. size and density). As shown in Figure 2A, a gradient of density media is created in a centrifuge tube. When a sample is subjected to high centrifugal forces, EVs migrate through the gradient, separating based on their buoyant density. The purified EVs were collected and tested for purity by TEM imaging and SE-HPLC. As shown in Figure 2B and Figure 2C, few protein aggregates and debris can be observed in the TEM images of both purified HEK293 and MSC EV samples, indicating the absence of large impurities. As shown in Figure 2D, in SE-HPLC chromatogram, the peak area of the main peak is over 99% of the total peak area, indicating the absence of small impurities. These results of the purity assays suggest that the purified EV samples are appropriate to be used as the standard sample to calibration the ELISA assay.

**Figure 2.**
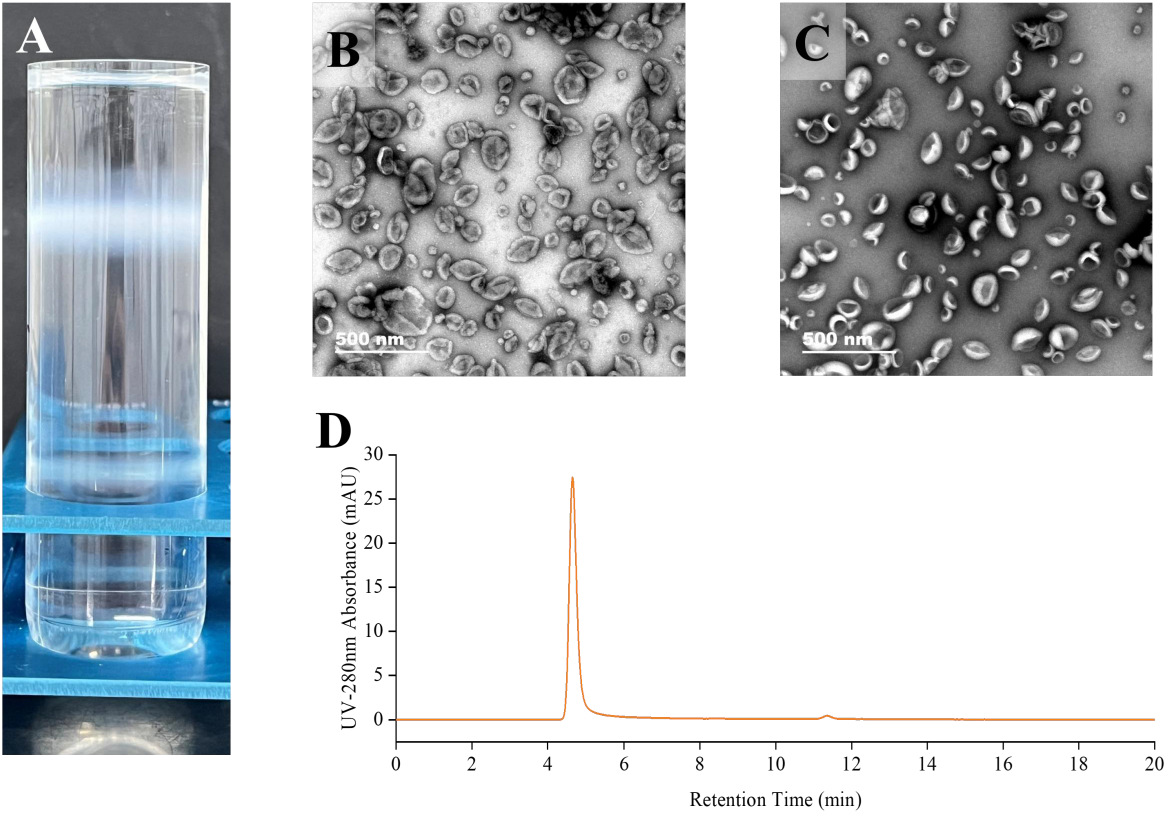
EV Standard used in the Vesicure total human EVs quantitation ELISA (CLIA) kit (A) Photo of Iodixanol density gradient ultracentrifugation purification (B) TEM image of the UGUC-purified HEK293 EV (C) TEM image of the UGUC-purified MSC-EV (D) SEC-HPLC chromatogram of the purified HEK293 EV

To establish the analytical performance of the Vesicure total human EVs quantitation ELISA (CLIA) kit, a series of experiments were performed to validate the specificity, accuracy, precision, detection range, and sensitivity, respectively.

A calibration curve was prepared by using HEK293 EV in the concentration range of 4.1E7 to 3E10 EVs/mL. As shown in Figure 3A, four-parameter-logistic (4PL) regression was applied to prepare the standard curve, and R^2^=0.9998 was obtained. The limit-of-detection (LOD) is defined as the concentration of the standard corresponding to the mean value of the background plus three times the standard deviation. The background values were measured in six replicates, and the mean and standard deviation were calculated. Interpolated with the fitting curve, the detection limit was determined to be 1.04E7 EVs/mL. Similarly, the limit-of-quantification (LOQ) is defined as the concentration of the standard corresponding to the mean value of the background plus ten times the standard deviation. And the quantification limit was determined to be 3.21E7 EVs/mL.

**Figure 3.**
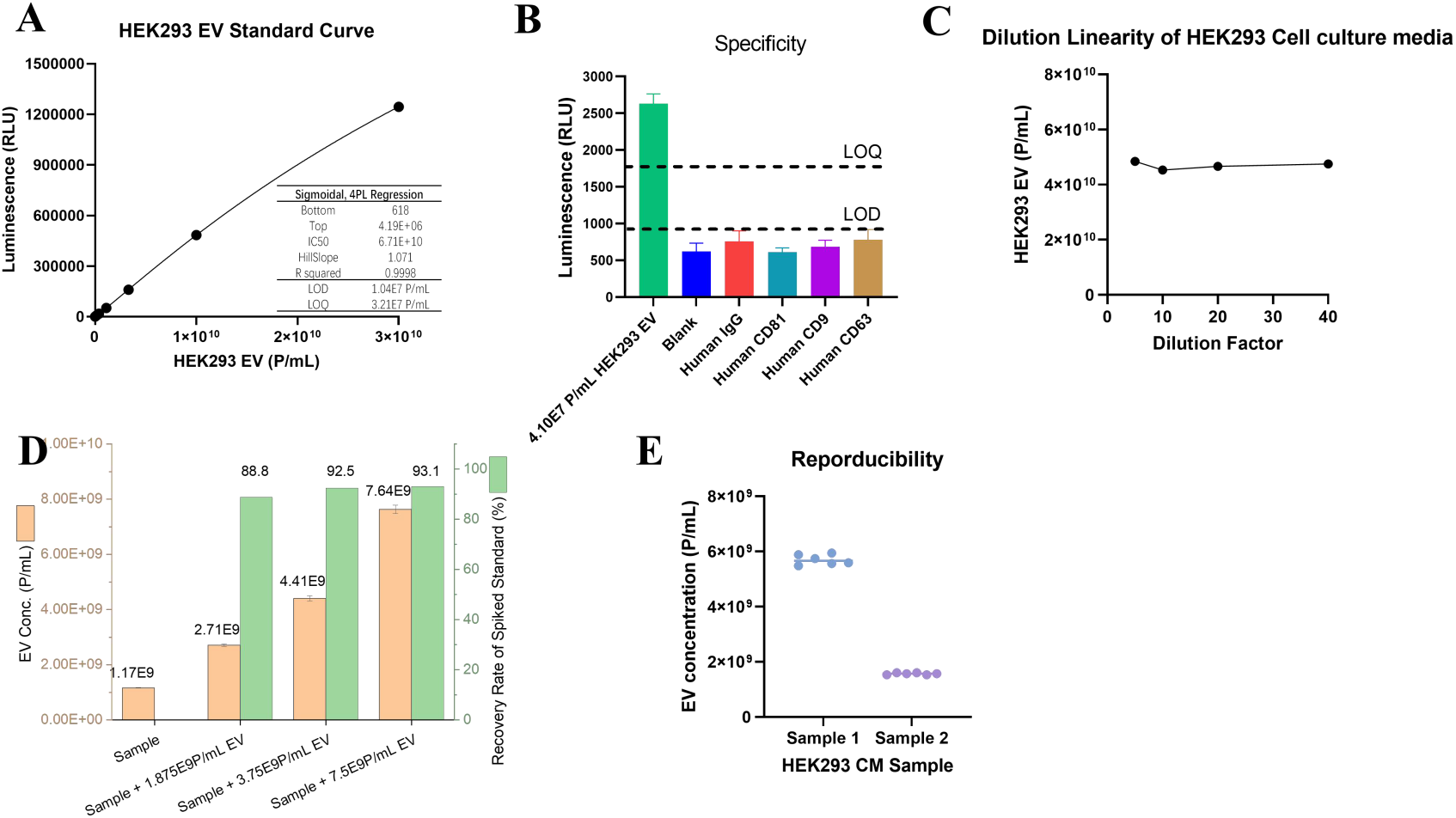
Validation of the Vesicure total human EVs quantitation ELISA (CLIA) kit (A) Standard Cruve (B) Specificity (C) Dilution Linearity (D) Recovery Rate of Spiked Standard (E) Reproducibility

The specificity of the ELISA assay is crucial for ensuring accurate and reliable results. Specificity refers to the assay’s ability to detect the EV without cross-reacting with non-EV impurities in the sample. As shown in Figure 3B, a positive control (4.10E7 particles/mL HEK293 EV), and non-EV samples, including Human IgG (10 ng/mL), Human CD81 (10 ng/mL), Human CD9 (1 ng/mL), Human CD63 (1 ng/mL), were tested with our ELISA kit, respectively. All the four non-EV samples were tested below the detection limit. This result suggests that the Vesicure total human EVs quantitation ELISA (CLIA) kit enjoys fairly good specificity.

The accuracy was determined by the dilution linearity and the recovery rate of spiked standard samples in of biological sample. As shown in Figure 3C, a Cell culture media sample of the HEK293 cell line samples were diluted with dilution factors of 8, 16, 32, 64-fold, respectively. The interpolated EV concentration of the original sample stayed constant, indicating minimum impact of matrix effect on the ELISA assay. As shown in Figure 3D, Standard EV sample with different concentrations were spiked into the 25-fold diluted HEK293 cell culture media sample. The recovery rate of spiked samples was in the range of 75%-125%. These results suggest that the Vesicure total human EVs quantitation ELISA (CLIA) kit enjoys good accuracy.

Precision is crucial in the EV quantification ELISA assay because it ensures the consistency and reproducibility of results across multiple tests or experiments. As shown in Figure 3E, to establish the precision of the Vesicure total human EVs quantitation ELISA (CLIA) kit, two different HEK293 cell culture media samples were tested for 6 replicates, respectively. For each sample, the coefficient of variation (CV) is less than 15%, indicating good precision of the assay.

The harvested cell culture media (CM) is the product of the upstream cell culture process in large-scale EV production, and the concentration of EV particles plays a crucial role in determining the final EV yield, making it an important indicator for optimizing cell culture processes. However, CM contains many impurities, and the most commonly used method for EV quantification, NTA, is interfered with by these impurities, preventing accurate measurement of EVs. As a proof of concept, we applied the Vesicure total human EVs quantitation ELISA (CLIA) kit on the EV quantification of MSC CM for upstream process development. As shown in Figure 4, eight samples of MSC cell culture media were collected for EV quantification, tested by ELISA, NTA, and TEM imaging. For ELISA, the crude samples were tested directly with MSC-EV standard sample that was shown in Figure 2C. For NTA and TEM, an extra ultracentrifugation purification step was performed prior to the measurement to ensure a solid readout. As shown in Figure 4A, after UC treatment of eight different CM samples, NTA was used to quantify the EV particle numbers. However, in the TEM images of the UC-purified samples shown in Figure 4B, it can be observed that although UC effectively purified and enriched the EVs from the CM, a numbers of protein aggregates and impurities still remained. Furthermore, multiple studies have shown that the recovery rate of UC-purified EVs is only around 20%, which further affects the accuracy of this quantification method. These data indicate that UC-NTA results are not sufficient for accurate EV quantification.

**Figure 4.**
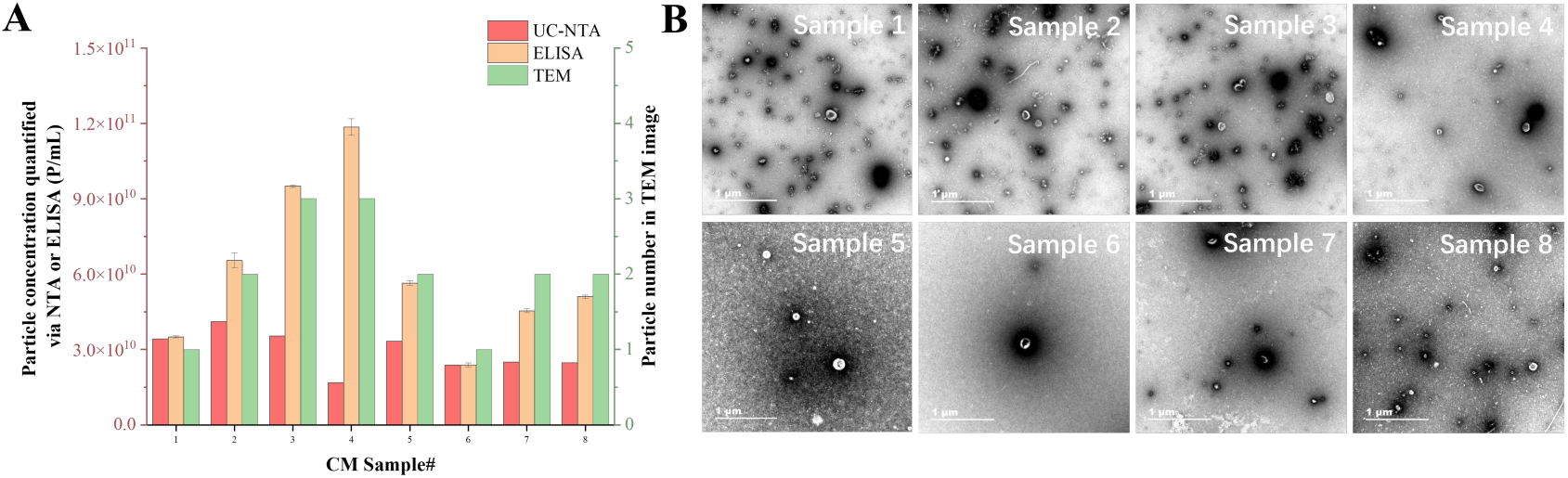
Application of the Vesicure human EV quantitation ELISA kit in Upstream process development (A) Comparison among ELISA, NTA, and TEM on different harvested cell-culture fluid samples (B) TEM images of UC-purified harvested cell-culture fluid samples

The sandwich ELISA method, by utilizing antibody-antigen binding, can specifically detect the EV concentration in crude samples. Testing showed that the recovery rate of spiked EVs in a 10-fold diluted MSC CM was 79.2%, meeting the requirement of 75-125%, indicating that this method can accurately quantify EV concentrations in the supernatant. Although TEM cannot be used for EV quantification, the number of EVs observed in the field of view can provide indirect insights into the EV concentration in the sample. For example, as shown in Figure 5B, despite the presence of impurities in samples 1, 2, and 3, the number of EVs showed an upward trend; in sample 4, most particles observed were EVs, and the EV count was the highest among all samples; in samples 5, 6, and 7, sample 6 was the purest but had the lowest EV count. By comparing the quantification results from ELISA with the number of EV particles observed in the TEM images of different samples, we found that, as shown in Figure 5A, the trend is highly consistent. This result further confirms the ability of the ELISA quantification kit to accurately quantify MSC EVs in the MSC cell culture supernatant.

**Figure 5.**
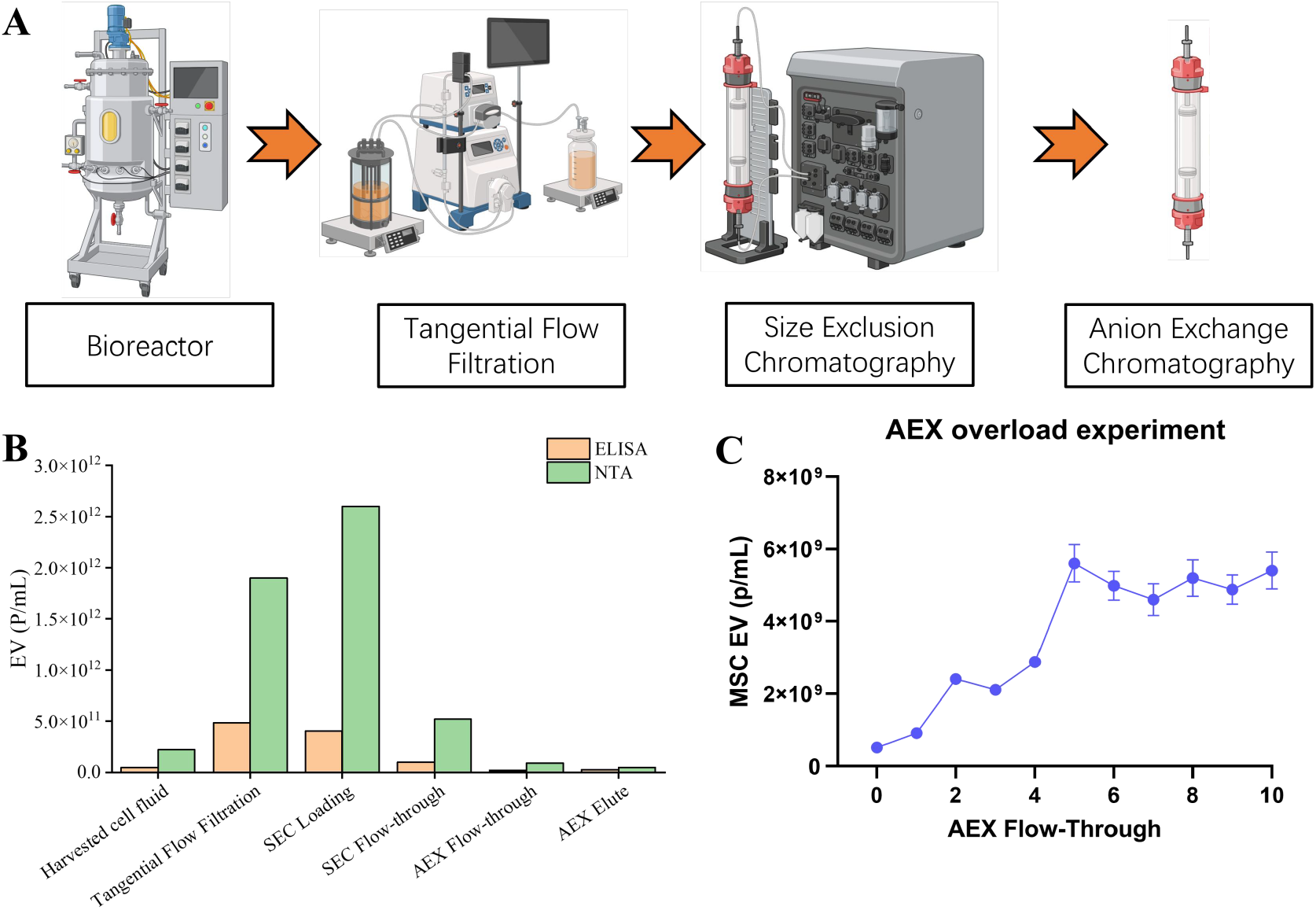
Application of the Vesicure human EV quantitation ELISA kit in Downstream process development (A) Workflow of the Vesicure Therapeutic Large-scale HEK293 EV purification process (B) Comparison between ELISA and NTA on EV samples through EV purification process (C) EV quantification by ELISA in AEX overload experiment for the purification EV.

Downstream process development is critical in large-scale EV manufacturing to ensure the efficient purification and isolation of high-quality EVs from complex mixtures, such as cell culture supernatant or human body fluids. This process involves removing impurities like proteins, cell debris, and other contaminants while maintaining the integrity and functionality of EVs. Key steps like filtration, chromatography, and ultracentrifugation must be optimized to maximize yield, purity, and consistency. As reported previously,^52^ we have developed a multi-step chromatography to achieve large-scale production of HEK293-derived EVs. As shown in Figure 5A, this strategy involves several steps, and precise EV quantification is essential during method development to optimize various parameters. However, the presence of protein aggregates and impurities prevents NTA from providing accurate quantification. In addition, a variety of buffer systems with different pH and conductivity were involve intermediate samples in downstream purification process, causing potential challenges for precise EV quantification.

Using an example from our downstream multi-step chromatography purification intermediates (Table 1), we compared the data obtained from ELISA and NTA quantification. As shown in Table 1, ELISA consistently yielded HEK293 EV recovery rates of over 85% for all process intermediate samples. As illustrated in Figure 5B, the quantification results from NTA and ELISA show significant differences in the initial purification steps because NTA was affected by protein aggregates while ELISA has a higher specificity for EVs. However, as purification progresses and the EV sample purity increases, the results from NTA and ELISA become more aligned. Figure 5 shows another application of the ELISA-based EV quantification method, in which the ELISA method was applied to monitor the EV concentration in the flow-through of the AEX column in an overload experiment. In this experiment, exceed amount of crude EV sample was introduced into the AEX column, and the flow-through was collected in tubes. Each sample were tested with the ELISA kit for EV quantitation. As shown in Figure 5C, the concentration of EV kept increasing and then reach the plateau along the introduction of the EV crude sample, indicating the saturation time point. Thus the column capacity can be calculated based on the data provided by EV quantification ELISA method. These results demonstrate the reliable role of ELISA-based EV quantification in supporting process optimization and control in large-scale EV production.

**Table 1.** Vesicure Downstream Process intermediate products in HEK293 EV purification.

| Sample | Buffer | ELISA Spiked Standard<br>Sample Recovery Rate (%) |
| --- | --- | --- |
| Harvested CM | Cell culture media | 86.0 |
| TFF | HEPES | 85.2 |
| SEC loading | HEPES | 93.1 |
| SEC flow-through | HEPES | 91.1 |
| AEX<br>flow-through | HEPES, NaCl | 90.4 |
| AEX elute | HEPES, NaCl | 89.6 |

## Conclusion

We have successfully developed a human EV quantification ELISA kit to support both the upstream and downstream process development and in-process control for large-scale EV manufacture. The detection range is 4.1E7 to 3E10 EVs/mL, with a LOD of 1.04E7 EVs/mL and a LOQ of 3.21E7 EVs/mL. This assay was validated in the aspect of specificity, accuracy, precision, and sensitivity. In addition, this kit was applied to harvested cell culture fluid samples from upstream process, and the ELISA was proven to be reliable by comparing with UC-NTA and UC-TEM. The ELISA kit was also applied to samples from downstream process to demonstrate the capability of support the in-process control as well as method development for EV purification. We believe the Vesicure Total Human EVs Quantification ELISA (CLIA) kit will make magnificent contributions to large-scale EV manufacturing and holds great potential for a variety of industrial, research, and clinical applications.

